# Direct and Indirect Models for Protein Chemical Denaturation are Characteristic of Opposite Dynamic Properties

**DOI:** 10.1101/073346

**Authors:** Liangzhong Lim, Linlin Miao, Jianxing Song

## Abstract

Two major models, namely direct and indirect models, have been proposed for the protein chemical denaturation but it remains challenging to experimentally demonstrate and distinguish between them. Here, by use of CD and NMR spectroscopy, we succeeded in differentiating the effects on a small but well-folded protein WW4, of GdmCl and NaSCN at diluted concentrations (≤200 mM). Both denaturants up to 200 mM have no alternation of its average structure but do reduce its thermodynamic stability to different degrees. Despite acting as the stronger denaturant, GdmCl only weakly interacts with amide protons, while NaSCN shows extensive interactions with both hydrophobic side chains and amide protons. Although both denaturants show no significant perturbation on overall ps-ns backbone dynamics of WW4, GdmCl suppresses while NaSCN enhances its μs-ms backbone dynamics in a denaturant concentration dependent manner. Quantitative analysis reveals that although they dramatically raise exchange rates, GdmCl slightly increases while NaSCN reduces the population of the major conformational state. Our study represents the first report deciphering that GdmCl and NaSCN appear to destabilize a protein following two models respectively, which are characteristic of opposite μs-ms dynamics.

---

**Table of Contents.**
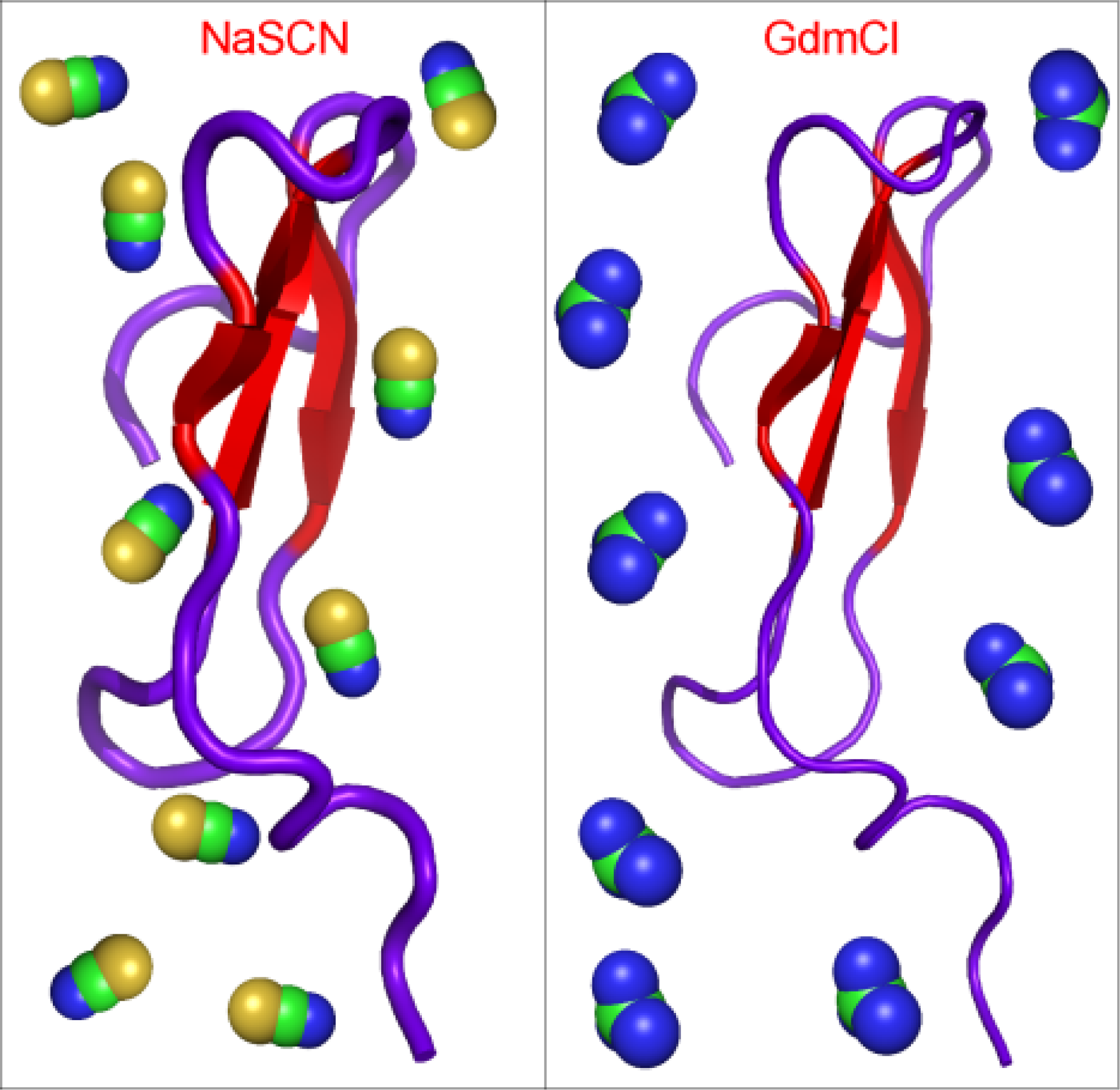
**NaSCN and GdmCl destabilize a protein with opposite dynamics:** Chemical denaturants are commonly utilized to study protein folding and stability but it still remains challenging to experimentally demonstrate and distinguish between direct and indirect models for their actions. Here NaSCN is revealed to destabilize WW4 by direct interactions while GdmCl by indirect effects. Most unexpectedly, two models are characteristic of opposite backbone dynamics. The study also suggests the absence of a simple correlation between protein dynamics and stability (see picture).

To function properly *in vivo*, many proteins fold into a specific three-dimensional structure called the native state which is only marginally stable relative to the denatured state. Thermodynamic stability is not only essential for biological functions of proteins, but also a key factor governing protein aggregation responsible for a variety of neurodegenerative diseases, as well as industrial applications. Consequently, the study of protein folding and stability has attracted the attention of protein chemists for about 100 years ^[1]^. One common method to experimentally investigate protein stability and folding process is involved in using chemical denaturants including guanidinium chloride (GdmCl) and sodium thiocyanate (NaSCN) which can modulate protein stability. However, despite extensive investigations with experimental and computational approaches, it still remains largely controversial for the microscopic mechanisms by which denaturants destabilize the native state of a protein ^[2-9]^. Currently two major models have been proposed, emphasizing either a direct interaction between protein and denaturants ^[4, 5]^, or an indirect effect mediated by the alteration of water structure ^[6, 7]^. Nevertheless, it has been challenging to experimentally demonstrate and distinguish between the two models because denaturants usually operate at the molar concentrations at which denaturant molecules are in close proximity to the protein chain ^[9]^.

Herein, we aimed to distinguish two models by differentiating effects of two denaturants, GdmCl and NaSCN at diluted concentrations (≤ 200 mM), on a 39-residue but well-folded WW4 domain derived from WWP1 protein. Previously we determined its NMR structure (2OP7) to adopt the classic WW fold, and also found no detectable aggregation even at a concentration up to 2 mM ^[10]^. Unlike a large and folded protein with most hydrophobic side chains buried in the core, WW4 is only composed of a flat three-stranded β-sheet whose hydrophilic and hydrophobic residues appear to be largely accessible, thus having advantage to sensitively sense effects of denaturant molecules. Using CD and NMR spectroscopy, here we characterized its conformations, stability, interactions and dynamics in the presence of two denaturants. The results suggest that two denaturants destabilize WW4 by following direct and indirect models, characteristic of opposite µs-ms dynamic properties.

We first assessed the conformations of WW4 at five different conditions, namely without denaturant, with GdmCl or NaSCN at 20 mM and 200 mM respectively (Figure 1a). The presence of the negative signal at 280 nm in near-UV CD spectra indicates that despite being small, WW4 does have a tight tertiary packing. Furthermore, very similar spectra strongly demonstrate that WW4 has no significant conformational change at five conditions. Nevertheless, two denaturants did trigger a reduction of stability. The melting temperature of WW4 without denaturant is ∼63.5°C, that reduced to ∼54.5 and ∼60.3°C respectively for those with 200 mM GdmCl and NaSCN (Figure 1b). GdmCl acts as a stronger denaturant than NaSCN, consistent with the classic ranking ^[2]^.

**Figure 1.**
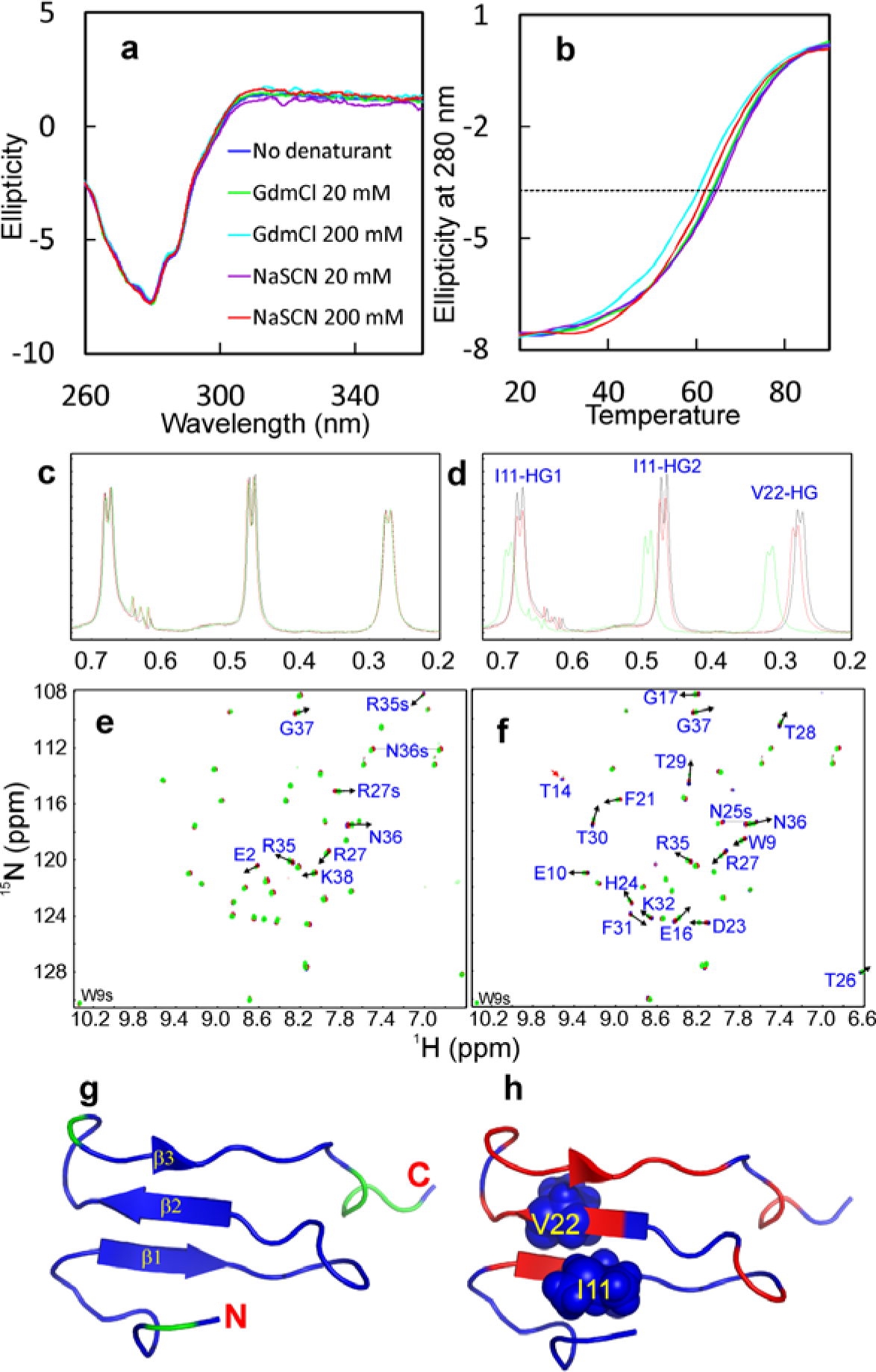
Effects on interaction, structure and stability. (a) Near-UV CD spectra of WW4 with GdmCl or NaSCN at different concentrations. (b) Thermal unfolding curves with GdmCl or NaSCN at different concentrations. One-dimensional ^1^H NMR spectra showing resonance peaks of Ile10 and Val22 methyl groups without (black) and with GdmCl (c) or NaSCN (d) at 20 mM (red) and 200 mM (green). Two-dimensional ^1^H-^15^N NMR HSQC spectra without (blue) and with GdmCl (e) or NaSCN (f) at 20 mM (red) and 200 mM (green). (g) WW4 structure with residues having significant shifts of backbone amide protons with 200 mM GdmCl colored in green. (h) WW4 structure with residues having significant shifts of backbone amide protons with 200 mM NaSCN colored in red. Ile10 and Val22 side chains are displayed in spheres.

Addition of GdmCl up to 200 mM triggered no significant change of resonance chemical shifts and shapes of representative Ile11 and Val22 methyl groups (Figure 1c), indicating no interaction with hydrophobic side chains. Further HSQC titrations show that GdmCl only weakly interacted with some well-exposed backbone and side chain amide protons (Figures 1e and 1g), consistent with previous results that GdmCl has no strong capacity to form hydrogen bonds with peptide groups ^[3]^, but weakly pair with positively charged groups ^[8]^. By contrast, addition of NaSCN led to a significant alternation of both chemical shift and line-broadening of methyl groups of Ile11 and Val22 (Figure 1d), which are located on the most structured first and second strands (Figure 1h). Moreover, NaSCN also triggers the shifts of many amide protons (Figure 1f), thus suggesting that NaSCN is able to extensively interact with both hydrophobic and charges groups. If considering such changes only occur at millimolar denaturant concentrations (mM), they are most likely resulting from direct contacts between denaturant molecules and affected protons ^[11, 12]^. Furthermore, HSQC spectral dispersions remain unchanged under five conditions, implying that no significant disruption of the tight packing occurs, consistent with the CD results.

To assess the dynamic effects of two denaturants, we acquired ^15^N backbone relaxation data including longitudinal relaxation time T1, transverse relaxation time T2 and {^1^H}-^15^N steady-state NOE (hNOE) for WW4 at five conditions (Figure S1). The similar hNOE values (Figure S1a) indicate that WW4 has no significant change in ps-ns backbone dynamics ^[13, 14]^. However, the differences were observed for T1/T2 values (Figure S1c), implying that WW4 has µs-ms dynamics under different conditions.

We then used “Model-free” formulism to analyze the data. First, we determined its overall rotational diffusion tensors (Table S1) by program ROTDIF ^[14]^. NaSCN induced the increase of overall rotational correlation time (Tauc) from 4.9 ns (no denaturant) to 5.4 (20 mM) and 6.1 (200 mM) ns respectively. Here, the value of TauC is expected to be mainly modulated by changes of the solution viscosity and alternation of WW4 dynamics upon adding denaturants. Surprisingly, although addition of GdmCl increases the solution viscosity, the apparent TauC values calculated from the relaxation data in fact decreased with the increase of GdmCl concentrations, 4.5 ns for 20 mM and 4.2 ns for 200 mM. This strongly implies that GdmCl is capable of reducing WW4 dynamics.

We subsequently conducted “Model-free” analysis by program Dynamics ^[14]^. Figure 2a presents the squared generalized order parameters, S^2^, of WW4 without denaturant, which reflect ps-ns conformational dynamics, ranging from 0 for high internal motion, to 1 for completely restricted motion. Excepted for the terminal residues Asn1-Leu5, Asn36-Ser39 and the loop residues Arg15-Glu16, the other non-Proline residues all have S^2^> 0.7 (Figures 2a and 2b), indicating that WW4 is well-folded. On the other hand, addition of denaturants indeed results in some alternations of the residue-specific S^2^ values slightly, particularly for those with 20 mM NaSCN (Figure 2c), implying the redistribution of ps-ns backbone dynamics. Nevertheless, average S^2^ values are very similar under all five conditions: 0.72 ± 0.01 (no denaturant); 0.71 ± 0.01 (20 mM NaSCN); 0.73 ± 0.01 (200 mM NaSCN); 0.71 ± 0.01 (20 mM GdmCl) and 0.73 ± 0.01 (20 mM GdmCl). The results suggest that addition of two denaturants even up to 200 mM does not significantly alter the overall backbone dynamics of WW4 on the ps-ns time scale.

**Figure 2.**
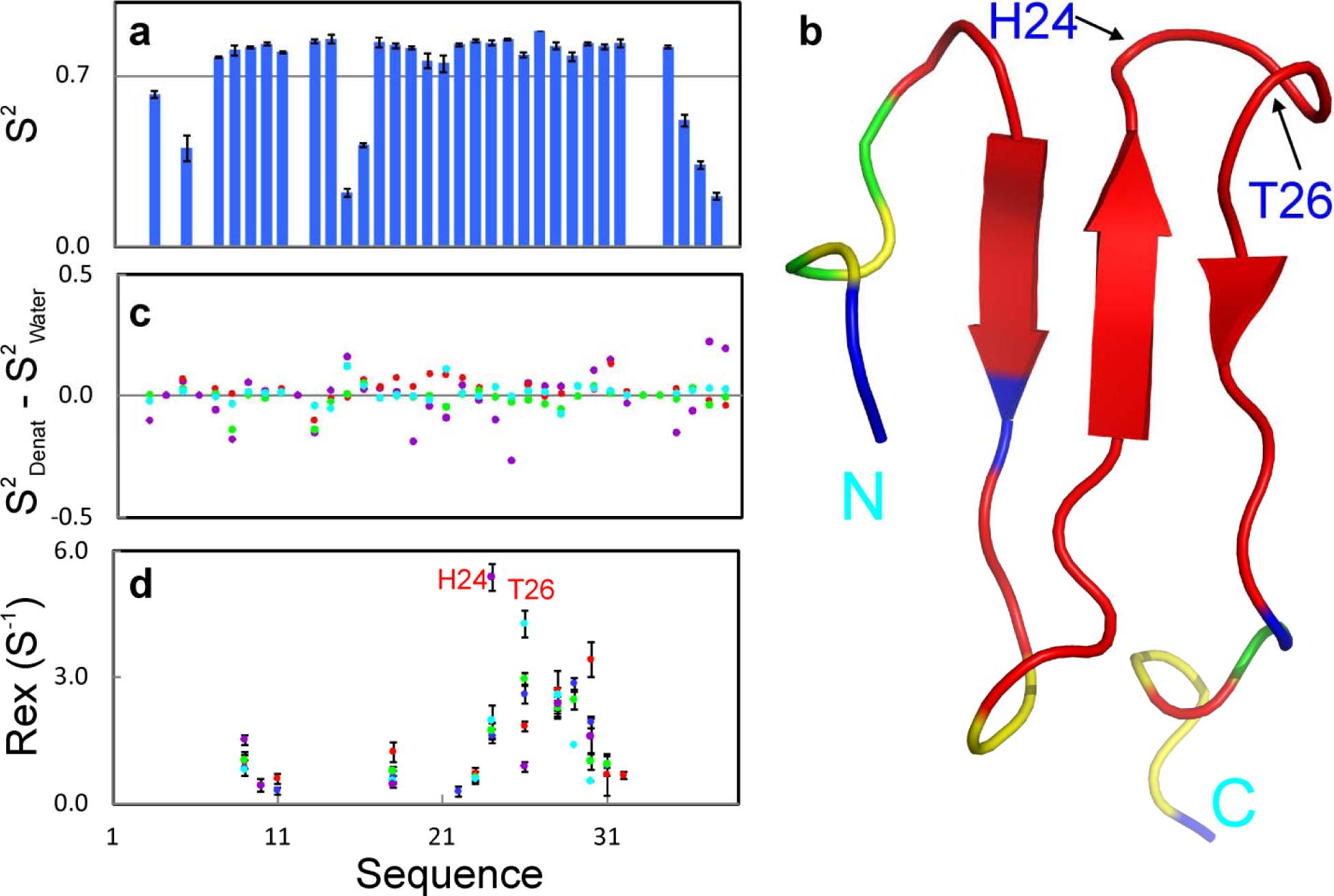
Model-free analysis. a) Squared generalized order parameters (S^2^) of WW4 without denaturant. b) WW4 structure colored based on S^2^ values of WW4 without denaturant: blue: absence of S^2^ due to the overlap of HSQC peaks and relaxation data of poor quality; green: Proline residues; red: S^2^ > 0.7 and yellow: S^2^ < 0.7. The locations of His24 and Thr26 are also indicated. c) Differences of S^2^ of WW4 with GdmCl at 20 mM (green), 200 mM (cyan); or NaSCN at 20 mM (purple); 200 mM (red) from those without denaturant. d) Residue-specific Rex of WW4 without (blue) and with GdmCl at 20 mM (green), 200 mM (cyan); or NaSCN at 20 mM (purple); 200 mM (red).

Model-free analysis also yields to Rex, which reflects conformational exchanges on the µs-ms time scale. As shown in Figure 2d, without denaturant, residues with Rex include Trp9, Ile11, Val18, Val22, Asp23, His24, Thr26, Thr28, Thr29, Thr30 and Phe31. In particular, His24, Thr26, Thr28, Thr29 and Thr30 have Rex > 1 Hz. Markedly, addition of two denaturants does not significantly change the overall patterns but does alter Rex value. For example, addition of GdmCl particularly at 200 mM significantly reduces Rex for most residues. By contrast, addition of NaSCN leads to the increase of Rex for most residues.

To independently confirm the effects of two denaturants on µs-ms backbone dynamics of WW4, we further performed ^15^N backbone CPMG relaxation dispersion measurements under five conditions (Figure 3). For WW4 without denaturant, the residues with ΔR_2_^eff^ > 4 Hz include Thr14, Gly17, Val18, Thr28, Thr29, Thr30 and Phe31. This pattern is consistent with that of Rex derived from Model-free analysis. Only His24 and Thr26 with large Rex show no significant dispersion response. One reasonable explanation is that the exchanging conformations involved in His24 and Thr26 may not have sufficient chemical shift difference. Consequently, their conformational exchanges could not be detectable by the CPMG relaxation dispersion measurements.

**Figure 3.**
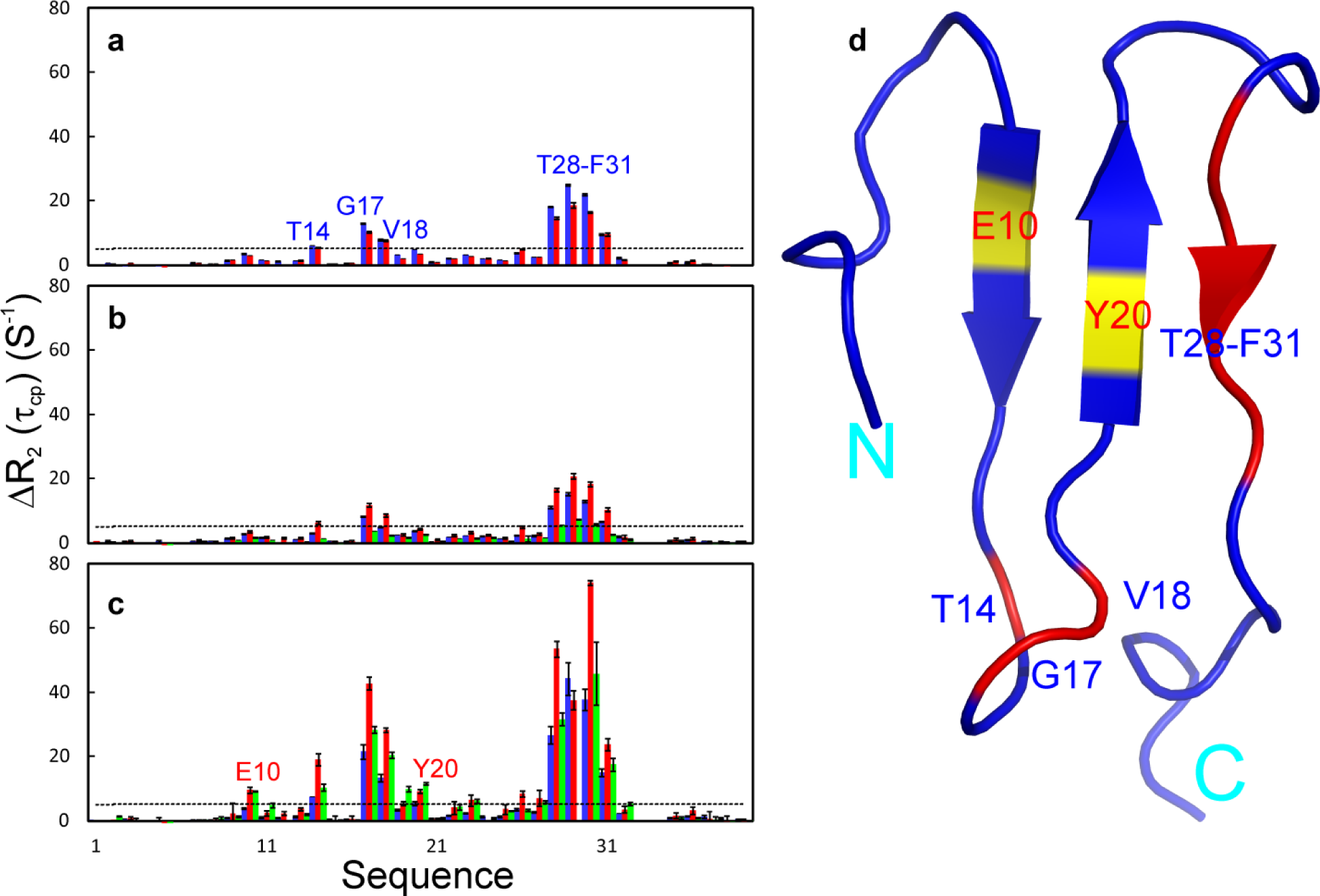
^15^N backbone CPMG relaxation dispersion. Differences of effective transverse relaxation rates (ΔR_2_^eff^) at 80 and 960 MHz, for WW4 without (a) and with GdmCl (b) or NaSCN (c). For WW4 without denaturant, blue bars are for data on 500 MHz while red on 800 MHz. For WW4 with 20 mM GdmCl or NaSCN, blue bars are for data on 500 MHz while red on 800 MHz. Green bars are for WW4 with 200 mM GdmCl or NaSCN on 500 MHz. d) WW4 structure colored based on ΔR_2_(τ_cp_) without denaturant, red is for residues with ΔR_2_(τ_cp_) > 4 Hz. Glu10 and Tyr20 are colored in yellow as their ΔR_2_(τ_cp_) became > 4 Hz upon adding 20 mM NaSCN.

Most strikingly, two denaturants oppositely mediate µs-ms conformational exchanges of WW4 in a concentration dependent manner. GdmCl suppresses whereas NaSCN enhances the exchanges for most residues (Figure 3 and Table S2). To gain a quantitative insight, by assuming a two-state conformational exchange, we fitted the data with ΔR^2eff^ values > 4.0 Hz on both 500 and 800 MHz by program GUARDD ^[15]^ to obtain detailed exchange parameters (Table S3). For WW4 without denaturant, the major conformational states have populations ≥ 90% for Thr14, Gly17, Val18, and Thr28-Phe31, which undergo exchanges with a minor state, with Rex ranging from 111.8 to 352.6 Hz. Interestingly, both denaturants significantly increase exchange rates Rex, and the increase is more significant for NaSCN than for GdmCl. Strikingly, GdmCl slightly reduces while NaSCN increases the population of the major state (Table S1).

As NaSCN has extensive interactions with both hydrophobic side chain and amide protons, and as such it appears to destabilize the protein by following the direct model. Very unexpectedly, although GdmCl acts as a stronger denaturant, it only weakly interacts with some well exposed amide protons. Therefore, GdmCl seemingly destabilizes the protein by following the indirect model. Here, it is also surprising to find that GdmCl and NaSCN behave so differentially in interacting with WW4. Previously, both guanidinium and thiocyanate ions were characterized to be weakly hydrated and in particular, guanidinium ion even has no recognizable hydration shell. Thus, both ions have been proposed to destabilize a protein by the preferential interactions with the protein surface ^[16]^.

Although GdmCl and NaSCN triggered no detectable alternation of the average conformation of WW4, they oppositely mediate µs-ms conformational exchanges in a concentration dependent manner. To the best of our knowledge, this represents the first uncovering that direct and indirect models for protein chemical denaturation are characteristic of the enhancement and reduction of µs-ms backbone dynamics. On the other hand, currently the relationship between protein dynamics and stability has been of fundamental interest but still remains controversial ^[17-22]^. Our results thus unambiguously indicate the absence of a simple correlation between them.

Mechanistically extensive interactions of NaSCN with both hydrophobic and hydrophilic groups appear to disrupt the tight packing, thus leading to the dynamic enhancement and reduction of the stability. It is so unexpected that GdmCl only weakly interacts with WW4, but can suppress µs-ms backbone dynamics and destabilize WW4 more. Here we propose that the presence of GdmCl may result in the alternation of water structure as extensively characterized [23]. This effect triggers the compression of WW4, which is characteristic of reduced µs-ms backbone dynamics. The compression may reduce or even remove the nano-cavity in the WW4 structure, which then leads to destabilization, as recently demonstrated for the pressure-triggered unfolding ^[24]^.

## Acknowledgments

We think Dr. Jing-song Fan for the assistance in acquiring NMR data. This study is supported by Ministry of Education of Singapore (MOE) Tier 2 Grant MOE2015-T2-1-111 to Jianxing Song. The funders had no role in study design, data collection and analysis, decision to publish, or preparation of the manuscript.

## Supporting Information

### Experimental Section

#### Materials and Methods

##### Expression and purification of WW4

The expression vector for WW4 was previously constructed (1). For bacterial expression of the recombinant proteins, the vector was transformed into *E. coli* BL21 cells. The cells were then cultured at 37 °C until the OD 600 value reached 0.6. Subsequently, IPTG was added into the broth to a final concentration of 0.3 mM to induce protein expression for 12 h at 20 °C. Cells were harvested by centrifugation and lysed by sonication in PBS buffer. The recombinant GST-fused WW4 protein was purified by affinity chromatography with glutathione-Sepharose 4B beads (Pharmacia Biotech) under native conditions. Subsequently, the WW4 domain was released from the GST fusion protein by on-gel thrombin cleavage at room temperature for 3 h, followed by further purification with HPLC on a reverse-phase C18 column (Vydac) eluted with the water-acetonitrile system by gradually increasing the acetonitrile concentration. To isotope label WW4 protein for ^1^H-^15^N NMR HSQC experiments, the recombinant protein was prepared with a similar protocol except the cells were grown in M9 medium with addition of (^15^NH4)_2_SO_4_ for ^15^N labeling.

##### Circular dichroism (CD) experiments

Since non-specific noise was extremely high over the far-UV region in the presence of GdmCl and NaSCN. In the present study, we characterized the structure and thermodynamic stability of WW4 by monitoring the near-UV region (260-360 nm). All near-UV CD spectra were collected at pH 6.4 in the absence and in the presence of two denaturants at different concentrations, on a Jasco J-810 spectropolarimeter equipped with a thermal controller as described previously (1, 2) with a protein concentration of 500 µM at 25 °C, using 1 mm path length cuvette with a 0.1 nm spectral resolution. Data from five independent scans were added and averaged. Thermal unfolding was performed on the same samples used for obtaining the above near-UV CD spectra with temperatures ranging from 20 to 90°C (3).

##### NMR titration of GdmCl and NaSCN to WW4

One WW4 stock sample was prepared by dissolving the protein powder in Milli-Q water (electric conductivity of 0.78 µS) to a final concentration of 250 µM and its pH was adjusted to 6.4 by adding diluted sodium hydroxide, which was subsequently split into individual NMR samples for titrations. The pH values of two denaturant solutions (1 M) were also adjusted to 6.4.

All NMR titration experiments were collected at 25°C on an 800 MHz Bruker Avance spectrometer equipped with a shielded cryoprobe as described previously (2). During titrations, series of one-dimensional ^1^H and two-dimensional ^1^H-^15^N HSQC spectra were acquired on the ^15^N-labeled WW4 domain at a concentration of 250 µM in the absence or in the presence of GdmCl and NaSCN at varying salt concentrations (3, 6, 10, 20, 30, 40, 60, 80, 100, 125, 150 and 200 mM) as we previously conducted with 14 salts (2). The pH values of the NMR samples for each titration were measured before and after titrations with 200 mM salts and found no detectable difference. NMR data were processed with NMRPipe (4) and subsequently analyzed with NMRView (5).

##### NMR characterization of ^15^N backbone dynamics on the ps-ns time scale

^15^N backbone T1 and T1ρ relaxation times and {^1^H}-^15^N steady state NOE intensities were collected for WW4 with a concentration of 500 µM at pH 6.4 under five conditions, namely WW4 without denaturant, with 20 mM GdmCl or NaSCN as well as with 200 mM GdmCl or NaSCN, on a Bruker DRX 500 MHz spectrometer equipped with pulse field gradient units at 25°C (6, 7). Relaxation time T1 was determined by collecting HSQC spectra with delays of 10, 80, 200, 320, 360, 420 and 500 ms using a recycle delay of 1 s, with a repeat at 200 ms. Relaxation time T1ρ was measured by collecting spectra with delays of 1, 30, 60, 90, 110, 130, 150 and 180 ms using a spin-lock power of 1.6 kHz and a 2.5 s recycle delay with a repeat at 90 ms. {^1^H}-^15^N steady-state NOEs were obtained by recording spectra with and without ^1^H presaturation, a duration of 3 s and a relaxation delay of 6 s.

Relaxation times were fitted to peak height data as single exponential decays. Spin-spin relaxation time T2 was calculated from T1ρ and T1 according to equation:

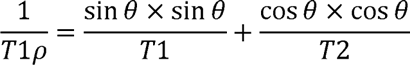

Where θ=atan(Δω/ω_1_) and Δω, ω_1_ are the resonance offset and spin-lock field strength, respectively (6).

NMR relaxation data were analyzed by “Model-Free” formulism with protein dynamics software suites (8, 9). Briefly, relaxation of protonated heteronuclei is dominated by the dipolar interaction with the directly attached ^1^H spin and by the chemical shift anisotropy mechanism (10-12). Relaxation parameters are given by:

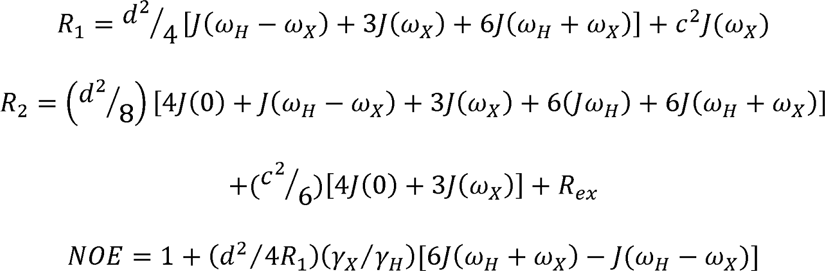

In which 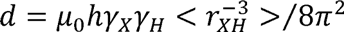, 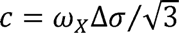, *μ*_0_ is the permeability of free space; *h* is Planck’s constant; *γ_X_γ_H_* are the gyromagnetic ratios of ^1^H and the X spin (X=^13^C or ^15^N) respectively; *γ_XH_* is the X-H bond length; *ω_H_* and *ω_H_* are the Larmor frequencies of ^1^H and X spins, respectively; and Δσ is the chemical shift anisotropy of the X spin.

The Model-Free formalism, as previously established (13) and further extended (12), determines the amplitudes and time scales of the intramolecular motions by modeling the spectral density function, *J*(ω), as

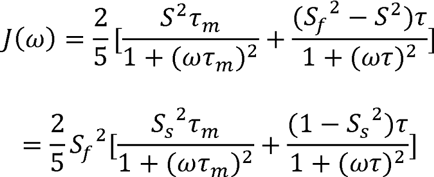

In which, *τ* = *τ_s_τ_m_*/(*τ_s_* + *τ_m_*), *τ_m_* is the isotropic rotational correlation time of the molecule, *τ_s_* is the effective correlation time for internal motions, 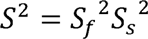 is the square of the generalized order parameter characterizing the amplitude of the internal motions, and 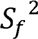 and 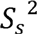 are the squares of the order parameters for the internal motions on the fast and slow time scales, respectively.

In order to allow for diverse protein dynamics, several forms of the spectral density function, based on various models of the local motion (11, 12, 14-16), were utilized, which include the original Lipari-Szabo approach (13), assuming fast local motion characterized by the parameters *S*^2^ and *τ_loc_*; extended model-free treatment, including both fast 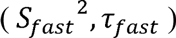 and slow 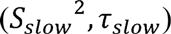 reorientations for the NH bond (*τ_fast_* ≪ *τ_slow_* < *τ_c_*); and could also allow for slow, micro- to milli-second dynamics resulting in a conformational exchange contribution, *R_ex_*. In the present study, the WW4 NMR structure (2OP7) with the lowest energy was used for “Model-free” analysis. For HSQC spectra of WW4 under five conditions, all peaks are well-separated and thus data are of high quality, except for the overlap of the Arg12 and Asp33 peaks (Fig. S1). Here, the overall rotational diffusion tensors and correlation time (TauC) of WW4 under five conditions were determined by ROTDIF (8), by selecting residues only with hNOE > 0.5 and having uniform T1/T2 values which include E3, L5, E7, G8, W9, E10, I11, Y13, R19, Y20, F21, V22, D23, N25, R27, K32 and R35. Subsequently “Model-free” analysis of relaxation data was performed by software DYNAMICS (9). We have analyzed the relaxation data with three overall models, namely isotropic, axially-symmetric and fully-anisotropic models and subsequently the axially-symmetric model was found to best describe WW4 under different conditions, whose overall rotational diffusion tensors and TauC are summarized in Table S1.

##### NMR characterization of ^15^N backbone dynamics on the µs-ms time scale

15N transverse relaxation dispersion experiments for WW4 with a concentration of 500 µM at pH 6.4 under five conditions were acquired on a DRX 500 and Bruker Avance 800 MHz spectrometers equipped with a z-axis gradient cryoprobe at 25°C (17). A constant time delay (*T*_CP_ = 50 ms) was used with a series of CPMG frequencies, ranging from 40, 80, 120, 160, 200, 240, 280, 320, 400, 480, 560, 640, 720, 800 and 960 Hz, with a repeat at 120 Hz. A reference spectrum without the CPMG block was acquired to calculate the effective transverse relaxation rate by the following equation:

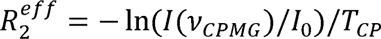

Where I(ν_CPMG_) is the peak intensity on the difference CPMG frequency, I_0_ is the peak intensity in the reference spectra.

Table S2 presents the difference of effective transverse relaxation rates (ΔR_2_^eff^) at 80 and 960 Hz for some WW4 residues under five conditions. The quality of the data of WW4 with 200 mM denaturants on the 800 MHz spectrometer is very poor as compared to the data on the 500 MHz spectrometer, mostly due to very weak or even disappeared peaks at the higher field (800 MHz) or/and high electronic noise. As a result, only the two-field (500 and 800 MHz) data for WW4 in the absence and in the presence of 20 mM GdmCl and NaSCN were analyzed by assuming a two-state conformational exchange (17-22), by the program GUARDD with the equation (21, 22):

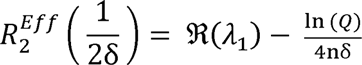

In which:

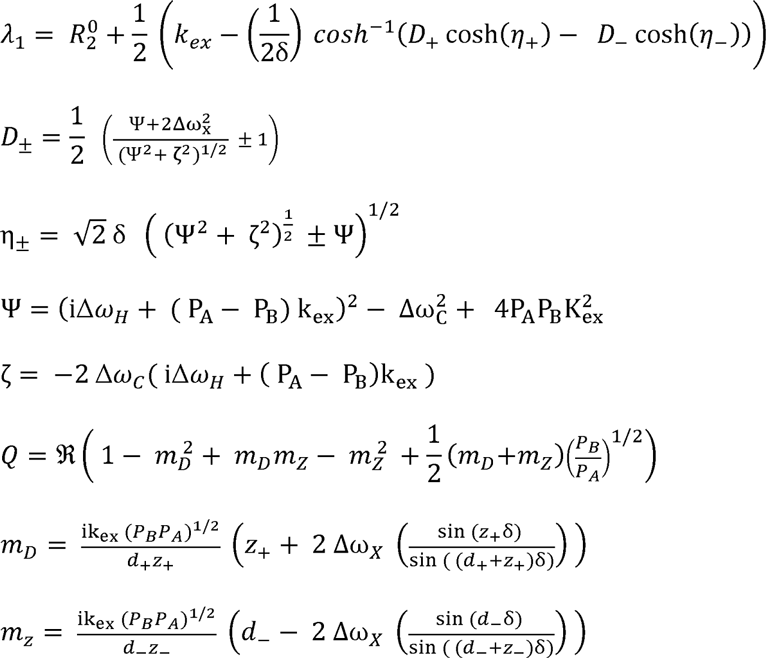

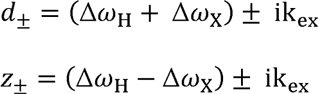

The obtained parameters defining conformational exchanges are presented in Table 1 and fitting curves directly output from the program GUARDD were shown in Figs. S2-S4.

## Supplementary Tables

**Table S1.** Diffusion tensors of WW4 derived from ^15^N relaxation data

|  | **Alpha** | **Beta** | **TauC** | **Dx=Dy** | **Dz** |
| --- | --- | --- | --- | --- | --- |
| **No denaturant** | 86 | 131 | 4.90 | $3.15 \cdot 10^7$ | $3.91 \cdot 10^7$ |
| **GdmCl (20 mM)** | 90 | 106 | 4.51 | $3.37 \cdot 10^7$ | $4.35 \cdot 10^7$ |
| **GdmCl (200 mM)** | 85 | 96 | 4.23 | $3.73 \cdot 10^7$ | $4.37 \cdot 10^7$ |
| **NaSCN (20 mM)** | 0 | 140 | 5.34 | $2.99 \cdot 10^7$ | $3.38 \cdot 10^7$ |
| **NaSCN (200 mM)** | 83 | 98 | 6.09 | $2.62 \cdot 10^7$ | $2.97 \cdot 10^7$ |

**Table S2.** Differences of effective transverse relaxation rates (ΔR_2_^eff^, Hz) for WW4 residues collected at 500 MHz

| Resid | $\Delta R_2^{\text{eff}}$ (Hz)<br>(No denaturant) | $\Delta R_2^{\text{eff}}$ (Hz)<br>(NaSCN 20 mM) | $\Delta R_2^{\text{eff}}$ (Hz)<br>(NaSCN 200 mM) | $\Delta R_2^{\text{eff}}$ (Hz)<br>(GdmCl 20 mM) | $\Delta R_2^{\text{eff}}$ (Hz)<br>(GdmCl 200 mM) |
| --- | --- | --- | --- | --- | --- |
| E10 | 3.6 | 3.9 | **9.6** | 2.8 | 1.7 |
| T14 | **6.0** | **7.5** | **18.9** | 3.1 | 1.3 |
| G17 | **12.9** | **21.5** | **42.8** | **8.2** | 3.7 |
| V18 | **8.0** | **13.4** | **28.3** | **4.9** | 2.3 |
| R19 | 3.2 | 3.4 | **5.4** | 2.4 | 1.7 |
| Y20 | **4.8** | **5.5** | **9.1** | 3.7 | 2.6 |
| V22 | 2.2 | 1.7 | **4.0** | 1.8 | 1.3 |
| D23 | 3.2 | 2.4 | **6.4** | 2.3 | 1.3 |
| T26 | 3.8 | 3.6 | **8.4** | 2.4 | 1.3 |
| R27 | 2.5 | 2.7 | **7.1** | 2.3 | 1.6 |
| T28 | **18.1** | **26.7** | **53.5** | **11.1** | **5.5** |
| T29 | **25.0** | **44.3** | **37.6** | **15.3** | **7.4** |
| T30 | **21.9** | **37.7** | **74.2** | **13.0** | **5.7** |
| F31 | **9.6** | **15.0** | **23.9** | **6.5** | 2.5 |

Note: the values > 4 Hz are bolded.

**Table S3.** Parameters Defining Conformational Exchanges

|  | Non |  |  | GdmCl<br>20 mM |  |  | NaSCN<br>20 mM |  |  |
| --- | --- | --- | --- | --- | --- | --- | --- | --- | --- |
| Residues | $\Delta\omega$<br>(ppm) | Pa<br>(%) | Rex<br>(Hz) | $\Delta\omega$<br>(ppm) | Pa<br>(%) | Rex<br>(Hz) | $\Delta\omega$<br>(ppm) | Pa<br>(%) | Rex<br>(Hz) |
| **E10** | | | | | | | $0.2 \pm 0.08$ | $98.1 \pm 13.2$ | $1287.0 \pm 74.0$ |
| **T14** | $0.25 \pm 0.02$ | $98.4 \pm 6.0$ | $352.6 \pm 149.1$ | $0.19 \pm 0.03$ | $98.6 \pm 0.3$ | $732.0 \pm 156.0$ | $0.16 \pm 0.03$ | $94.6 \pm 9.5$ | $904.7 \pm 177.9$ |
| **G17** | $0.33 \pm 0.02$ | $90.6 \pm 12.3$ | $111.8 \pm 107.8$ | $0.25 \pm 0.01$ | $97.6 \pm 0.2$ | $612.4 \pm 80.9$ | $0.25 \pm 0.01$ | $92.4 \pm 0.3$ | $918.1 \pm 50.1$ |
| **V18** | $0.28 \pm 0.02$ | $97.5 \pm 7.9$ | $308.9 \pm 129.2$ | $0.21 \pm 0.02$ | $98.2 \pm 0.2$ | $724.8 \pm 114.1$ | $0.19 \pm 0.01$ | $93.8 \pm 0.8$ | $923.0 \pm 85.6$ |
| **Y20** | | | | | | | $0.22 \pm 0.01$ | $98.1 \pm 0.1$ | $632 \pm 81.4$ |
| **T28** | $0.35 \pm 0.02$ | $92.5 \pm 9.6$ | $199.4 \pm 130.3$ | $0.27 \pm 0.15$ | $97.1 \pm 0.2$ | $819.6 \pm 87.9$ | $0.23 \pm 0.01$ | $89.9 \pm 0.8$ | $1064.2 \pm 78.0$ |
| **T29** | $0.44 \pm 0.03$ | $91.7 \pm 12.4$ | $254.4 \pm 169.6$ | $0.35 \pm 0.02$ | $96.9 \pm 0.2$ | $878.7 \pm 106.7$ | $0.34 \pm 0.04$ | $87.3 \pm 11.8$ | $422.2 \pm 216.1$ |
| **T30** | $0.33 \pm 0.02$ | $90.0 \pm 12.0$ | $166.1 \pm 118.6$ | $0.27 \pm 0.02$ | $96.6 \pm 0.2$ | $712.2 \pm 85.1$ | $0.25 \pm 0.01$ | $88.9 \pm 0.8$ | $967.2 \pm 83.2$ |
| **F31** | $0.25 \pm 0.02$ | $96.5 \pm 8.5$ | $269.3 \pm 110.8$ | $0.21 \pm 0.01$ | $97.8 \pm 0.2$ | $665.4 \pm 79.2$ | $0.24 \pm 0.02$ | $95.1 \pm 0.5$ | $571.1 \pm 105.0$ |

## Supplementary Figures

**Figure S1.**
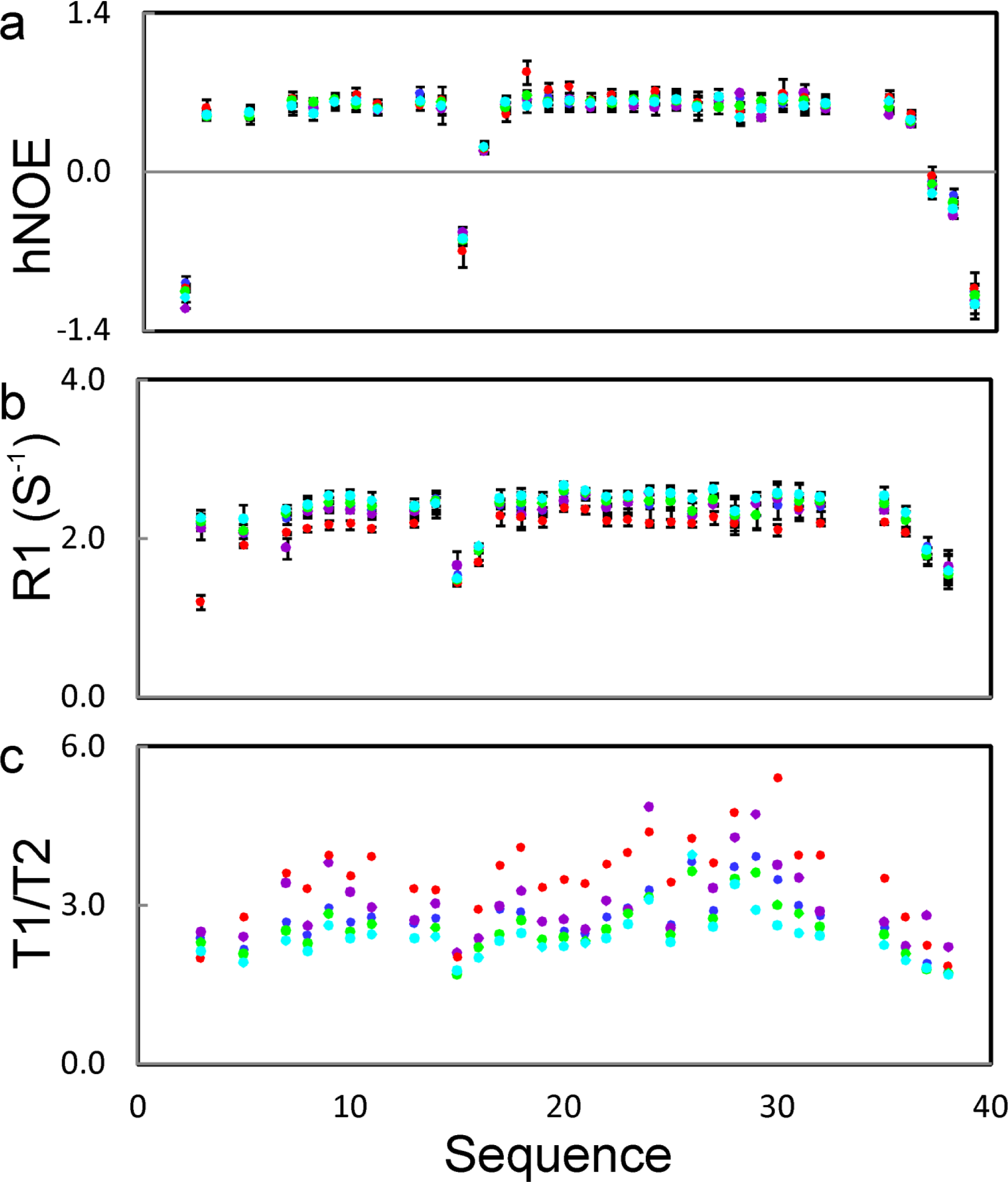
^15^N backbone relaxation data of WW4 under five conditions. a) {^1^H}-^15^N steady state NOE intensities (hNOE). b) Inverse of longitudinal relaxation time, T1 (R1). c) T1 divided by T2 (transverse relaxation time). Blue: WW4 without denaturant; green: with GdmCl at 20 mM; cyan: with GdmCl at 200 mM; purple: with NaSCN at 20 mM; red: with NaSCN at 200 mM.

**Figure S2.**
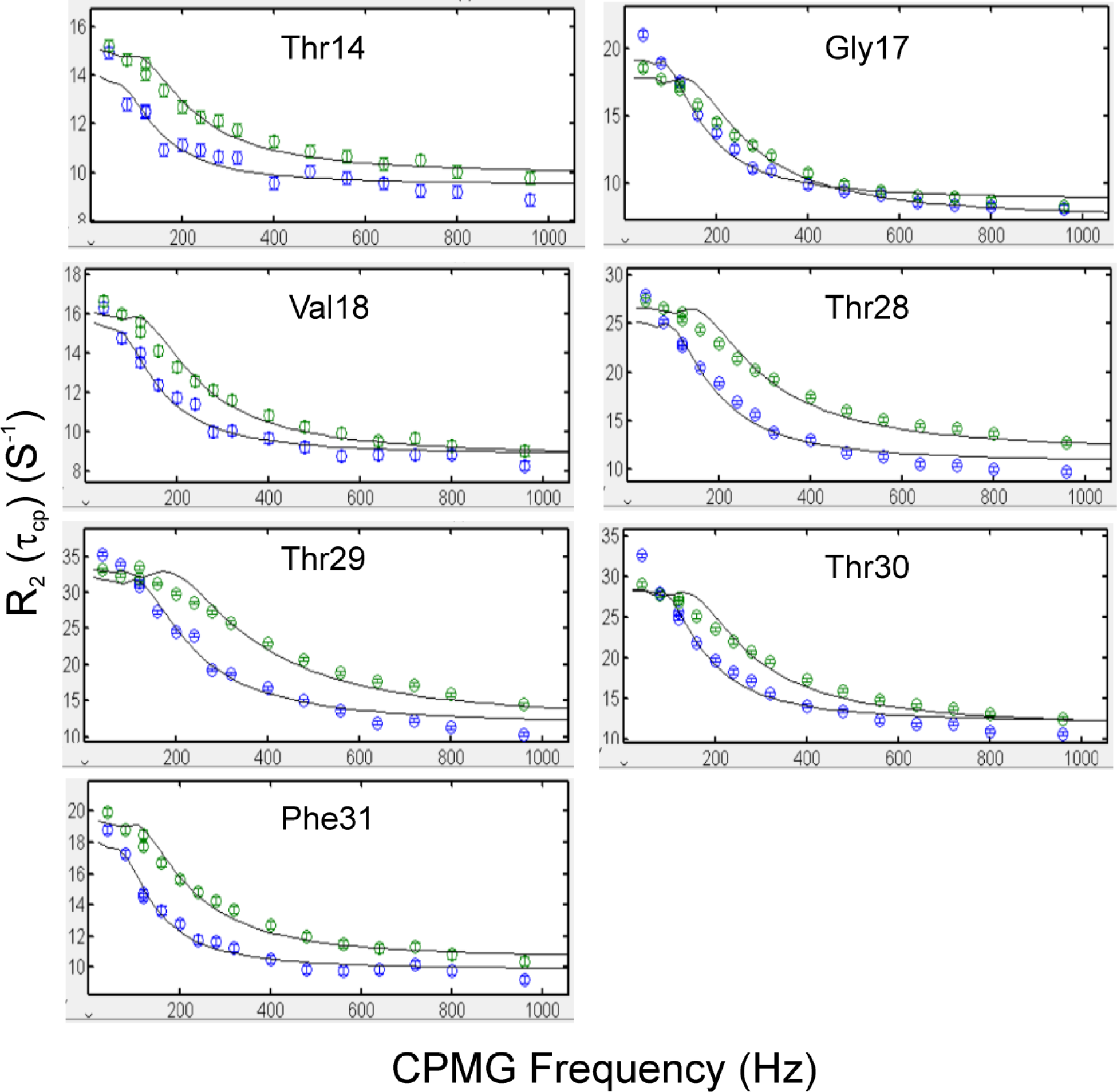
Fitting curves for WW4 without denaturant. Displayed are fitted (black lines) and experimental (circles) values of effective transverse relaxation rates (R_2_^eff^) for WW4 ^15^N backbone collected on 500 MHz (blue) and 800 MHz (green) spectrometers. X-axis is R_2_^eff^ (Hz) while y-axis is CPMG frequencies (Hz).

**Figure S3.**
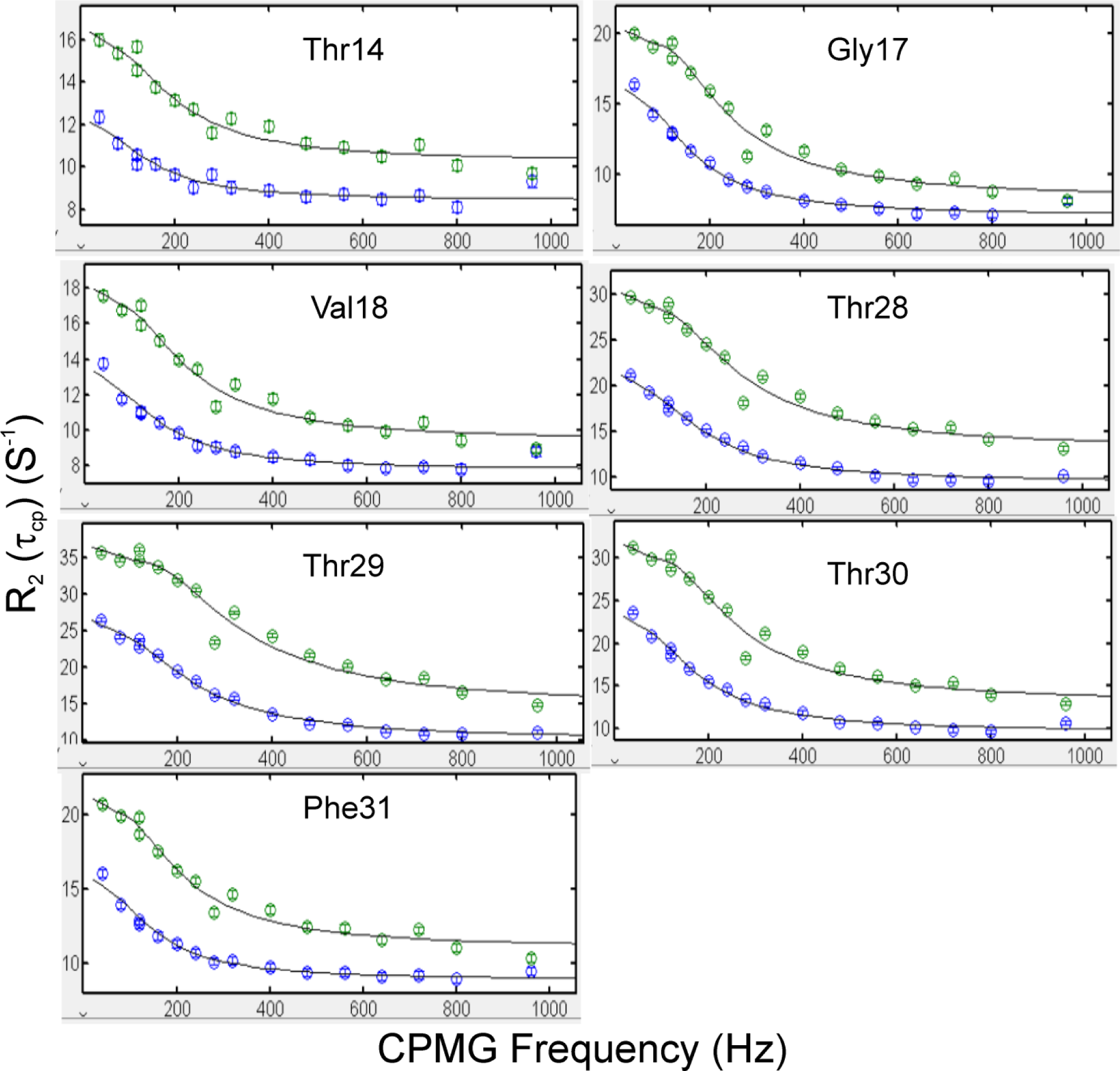
Fitting curves for WW4 with 20 mM GdmCl. Displayed are fitted (black lines) and experimental (circles) values of effective transverse relaxation rates (R_2_^eff^) for WW4 ^15^N backbone collected on 500 MHz (blue) and 800 MHz (green) spectrometers. X-axis is R_2_^eff^ (Hz) while y-axis is CPMG frequencies (Hz).

**Figure S4.**
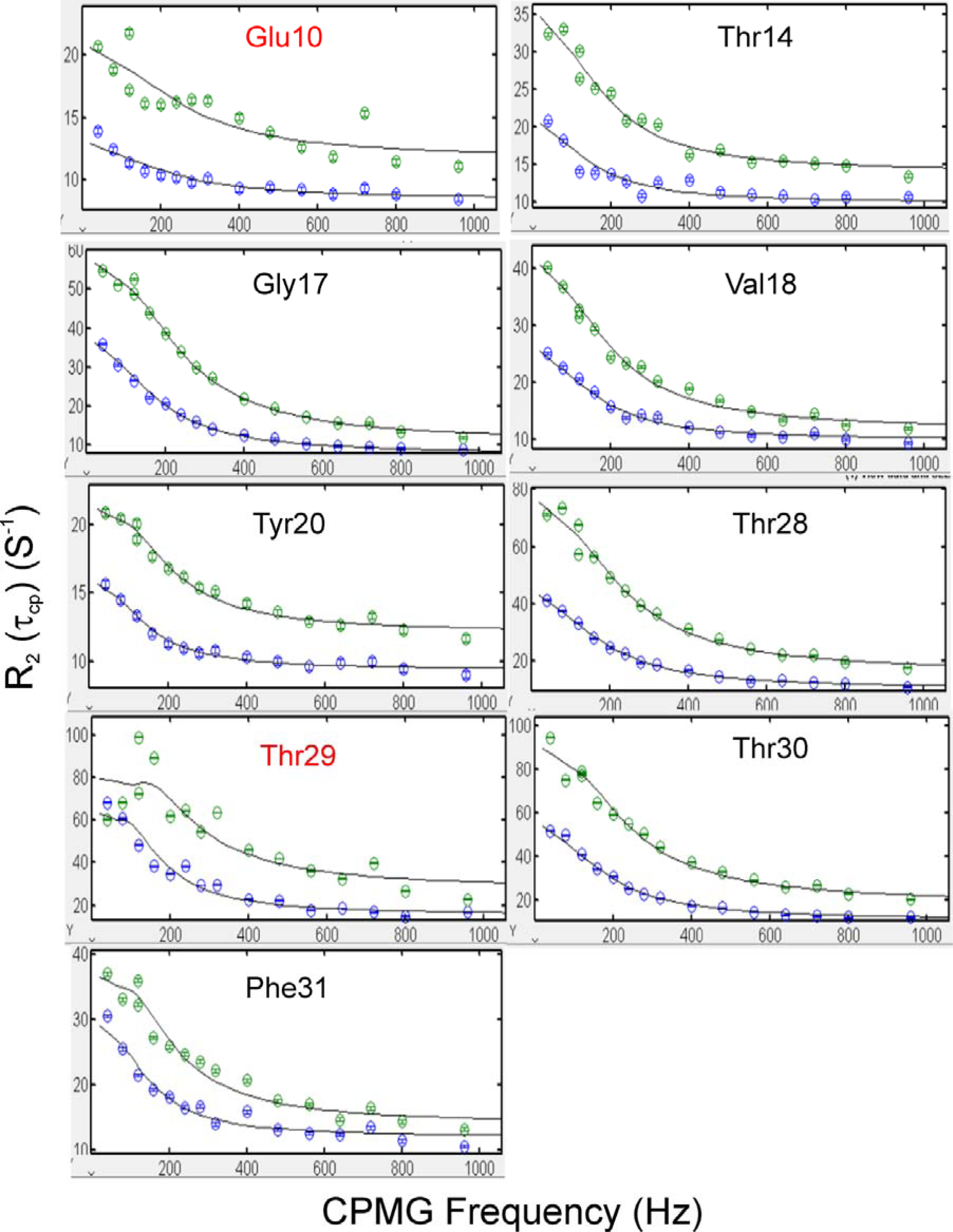
Fitting curves for WW4 with 20 mM NaSCN. Displayed are fitted (black lines) and experimental (circles) values of effective transverse relaxation rates (R_2_^eff^) for WW4 ^15^N backbone collected on 500 MHz (blue) and 800 MHz (green) spectrometers. X-axis is R_2_^eff^ (Hz) while y-axis is CPMG frequencies (Hz).

